# Comprehensive expression data for two honey bee species, *Apis mellifera* and *Apis cerana japonica*

**DOI:** 10.1101/2024.12.11.627317

**Authors:** Kakeru Yokoi, Masatsugu Hatakeyama, Seigo Kuwazaki, Taro Maeda, Mikio Yoshiyama, Mari Horigane-Ogihara, Shigeru Matsuyama, Akiya Jouraku, Hidemasa Bono, Kiyoshi Kimura

## Abstract

Comprehensive expression datasets were constructed for *Apis mellifera* and *Apis cerana japonica*. Post-oviposition day 6 to day 58 samples of *A. mellifera* workers (larva to adult); day 9, 10, 12, and 13 samples of *A. mellifera* queen (larva to pupa); and day 9 to day 18 samples of *A. cerana japonica* workers (larva to adult) were prepared, and RNA-Seq data were obtained. For *A. cerana japonica*, reference transcript sequence data, predicted amino acid sequence data, and functional annotation data were generated based on the genome sequence and RNA-Seq data. Using the transcript sequence and RNA-Seq data, comprehensive expression data for all transcripts of *A. mellifera and A. cerana japonica* were prepared. Hierarchical clustering analyses and the used sample preparation method ensured that both sets of expression data were reliable for use as comprehensive reference expression datasets. Therefore, these data are applicable for honey bee research or for comparative or evolutionary studies on insect species or social insect species at the genetic and molecular levels.

## Background

Honey bees (Hymenoptera: Apidae) are industrially important as producers of honey, royal jelly, propolis, and wax. Further, honey bees are pollinators and are essential for several crops (e.g., strawberry and watermelon). They are used in biological research as representative social insects^1^. A honey bee colony consists of a queen, workers (females), and drones (males). The queen and drones are mainly responsible for reproduction, whereas workers are mainly responsible for non-reproductive tasks, such as nursing larvae and foraging for nectar and pollen. There are several species of honey bees with species-specific traits.

The traits of *A. mellifera* include abundant honey production and easy rearing compared to other wild honey bee species. Moreover, *A. mellifera* has been used as a model social insect species to study social behavior, memory, and learning. Because of its importance, the first genome sequence data for *A. mellifera* were reported in 2006^2^. After several improvements, chromosome-level genome sequencing data were published^3^. Transcriptome analyses have also been performed using the genome sequence data for genetic or molecular level insights into *A. mellifera* or honey bee traits or reactions, such as casting or immune reactions against pathogens^4–7^, leading to the accumulation of *A. mellifera* RNA-Seq data in public databases. Using these RNA-Seq data, reference transcriptome data of *A. mellifera* with functional annotation data have been constructed^8^.

The Asian honey bee *Apis cerana*, a wild honey bee species distributed in Asia, is utilized for beekeeping in some rural areas^9^. The traits of *A. cerana* are more mildness, more frequently absconding and producing lower honey compared to *A. mellifera*. Furthermore, *A. cerana* show greater resistance to varroa mite and American foulbrood. The Japanese honey bee, *A. cerana japonica*, is a subspecies of *A. cerana*, which inhabits all areas of Japan except Okinawa and Hokkaido^10^. *A. cerana japonica* was used for beekeeping in Japan before *A. mellifera* were introduced. One interesting and characteristic trait of *A. cerana japonica* is the formation of a hot defensive bee ball against colony invaders such as the Japanese giant hornet *Vespa mandarinia*^11,12^. To advance research using *A. cerana* at the genetic and molecular levels, genome sequence data of *Apis cerana cerana* (Korean and Chinese strains) were published in 2015 and 2018, respectively, and chromosome-level genome sequence data were published in 2020^13–15^. The draft genome data of *A. cerana japonica* were reported in 2018, and chromosome-level genome data, constructed using *A. cerana* chromosome genome data, were published in 2023^16,17^. An RNA-Seq analysis of *A. cerana* has been performed. For instance, genes associated with hot defensive bee balls have been identified using RNA-Seq analysis^18^. Additional RNA-Seq analyses will be useful for *A. cerana* research. Notably, remarkable differences in traits are observed between *A. cerana japonica* and *A. mellifera* even though the two species are evolutionarily very close to each other.

Considering the available genome and RNA-Seq data for *A. mellifera* and *A. cerana japonica*, reference transcriptome data (expression data) of intact honey bees at different developmental stages are required. These data are useful for honey bee research at the genetic and molecular levels considering several publications on *A. mellifera* and *A. cerana japonica* using RNA-Seq or genome sequence data, as described above. However, preparing samples under completely unified conditions, which is relatively easy for model insects such as *Drosophila melanogaster* and *Bombyx mori*, is a very difficult task for honey bees, because of several reasons. First, honey bees cannot be reared from eggs to adults under artificial conditions such as in the laboratory. Honey bee larvae must be fed by nursing workers in the colony. Thus, the timing of feeding and conditions vary for each brood. Another factor is environmental conditions. Colonies are known to be placed in outside. Therefore, the temperature and weather conditions in which the colonies are placed are not stable, which affects the length of each stages^1^. The process of honey bee reproduction is another challenge. Queens mate with drones from their own and other colonies to oviposit the eggs of workers or new queens, leading to genetic variations in broods born from one queen. Furthermore, rearing *A. cerana japonica* is rather difficult because of its frequent absconding behavior. Nevertheless, comprehensive expression data for honey bees at multiple developmental stages under unified conditions are desirable. To the best of our knowledge, such a series of comprehensive expression data has not been previously published. These expression data are required for understanding honey bees and for entomological research communities. In this study, we prepared RNA-Seq data of intact two honey bee species (*A. mellifera* and *A. cerana japonica*) at multiple developmental stages using a sampling method with as little variation as possible (Sample preparation is described in the “Methods” section). Using RNA-Seq data, comprehensive expression data of prepared samples were constructed (Figure 1).

**Fig. 1.**
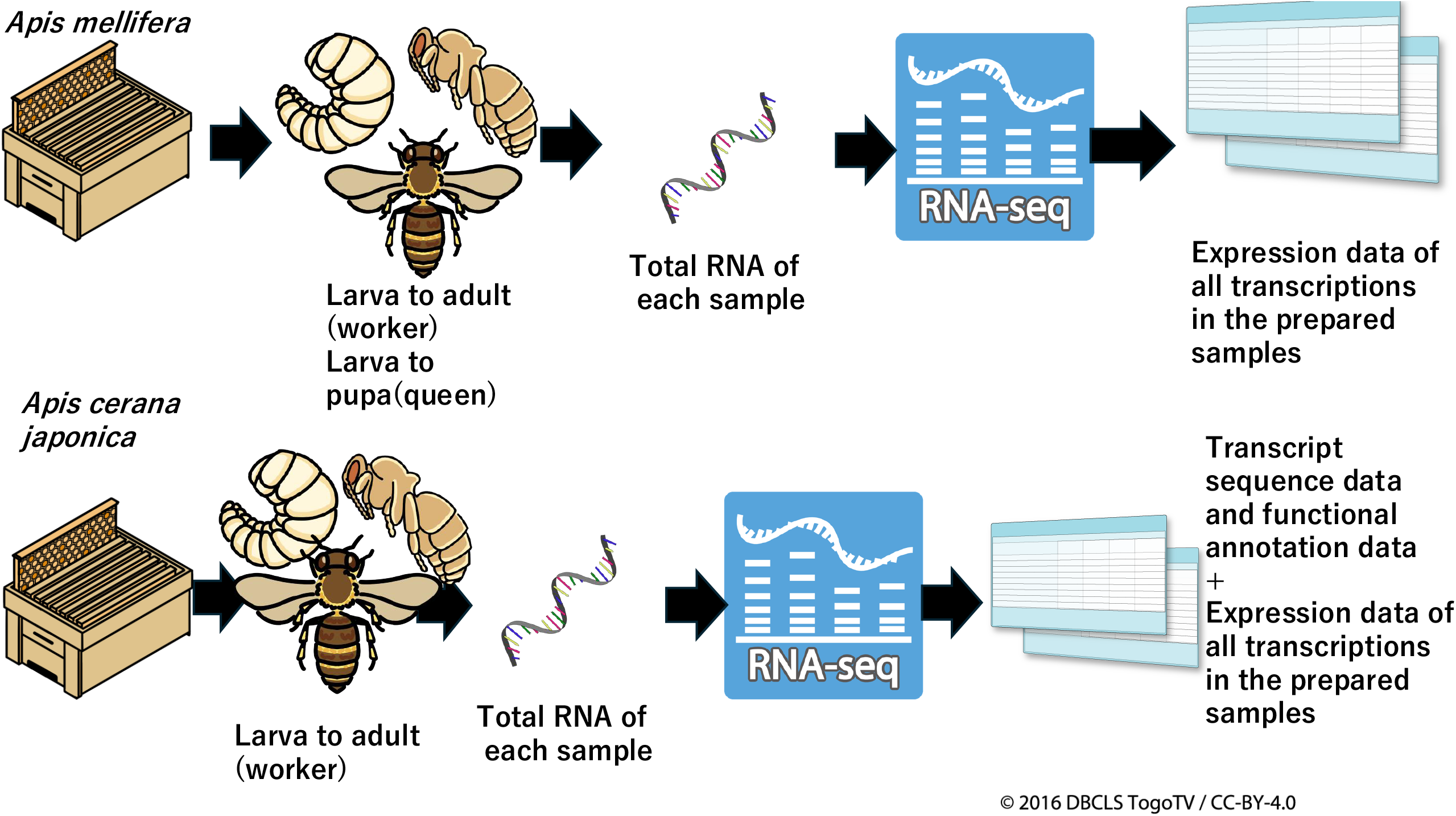
Schematic diagram of the study. A schematic representation of the study is presented. For *A. mellifera*, worker samples from larva to adult and queen samples from larvae to pupae were prepared. Total RNA was extracted from the samples and used for RNA-Seq. Using RNA-Seq data, expression data for all previously prepared transcripts in the prepared samples were constructed. For *A. cerana japonica*, larva to adult worker samples were prepared. Using the RNA-seq data, transcript sequence data, predicted amino acid sequence data, and functional annotation data were constructed. Furthermore, the expression data of the transcripts in the prepared samples were constructed using RNA-Seq and transcript sequence data.

## Materials and Methods

### Honey bee rearing

*A. mellifera* used in this study were from an Italian hybrid strain reared in a National Agricultural Research Organization (NARO) apiary; the bees were reared according to standard beekeeping practices. A healthy queen was used for sampling from a randomly selected hive.

For sampling *A. cerana japonica*, a swarm colony was reared in an apiary on the campus of the University of Tsukuba the year before sampling.

### Sample preparation

Samples for RNA-Seq, which was conducted from June to August 2021, were prepared by using an egg-laying box of the special queen rearing equipment from Ezi Queen Systems (Ezi Queen Technology Limited, Auckland, New Zealand), from in which a queen could freely lay eggs without escaping, and in which only workers could pass through the cover slits and care for the eggs and broods in the box (Fig. 2). Hereafter, “egg-laying boxes of the special queen rearing equipment from Ezi Queen Systems” will be referred to as “egg-laying boxes”. A single queen of both *A. mellifera* and *A. cerana japonica* was kept in the egg-laying box from 10 AM, and after 6 hours, the queen was released from the box. The oviposited eggs were used for RNA-seq analysis. At 10 AM after five days, sampling was started, and the obtained samples were stored at -80 °C until used. Similarly, other samples were consistently prepared at 10 AM. As shown in Fig. 3, the *A. mellifera* worker day 6 samples (AmW_d6) were samples obtained at 138-144 hours post-oviposition. Similarly, the postoviposition time of each sample can be determined by the sample name (Fig. 3). After sealing all the brood cells in the boxes (occurring in approximately day 8 to day 9 workers^1^), they were kept in incubators controlled at 35°C. For adult samples, emerged worker bees were marked on their backs and returned to their original hive within 24 h. The age of the worker bees was determined in this manner. Sampling continued in this way until sufficient numbers were obtained. The same queen was used for sampling. The nomenclature for each sample is shown at the bottom of Fig. 3. For example, AmW_d14_2 denotes “*A. mellifera* worker day 14, biological replicate 2” sample. *A. mellifera* worker samples were prepared from day 6 (larvae) until day 58 (adults), whereas *A. mellifera* queen samples were prepared from day 9 (larvae) until day 13 (pupae). *A. cerana japonica* worker samples were prepared from day nine (larvae or pre-pupa) until day 18 (adults) (Fig. 4). The biological replicates of these samples varied from one to five, and biological replicate numbers of some categories did not begin with “1” and were not sequential because sufficient amounts of total RNA for RNA-Seq were not extracted from some samples. Furthermore, *A. mellifera* queen day 11 and *A. cerana japonica* worker day 11 samples were excluded from the sample collections because there were no sample from which sufficient amount and quality of total RNA for RNA-Seq were extracted.

**Fig. 2.**
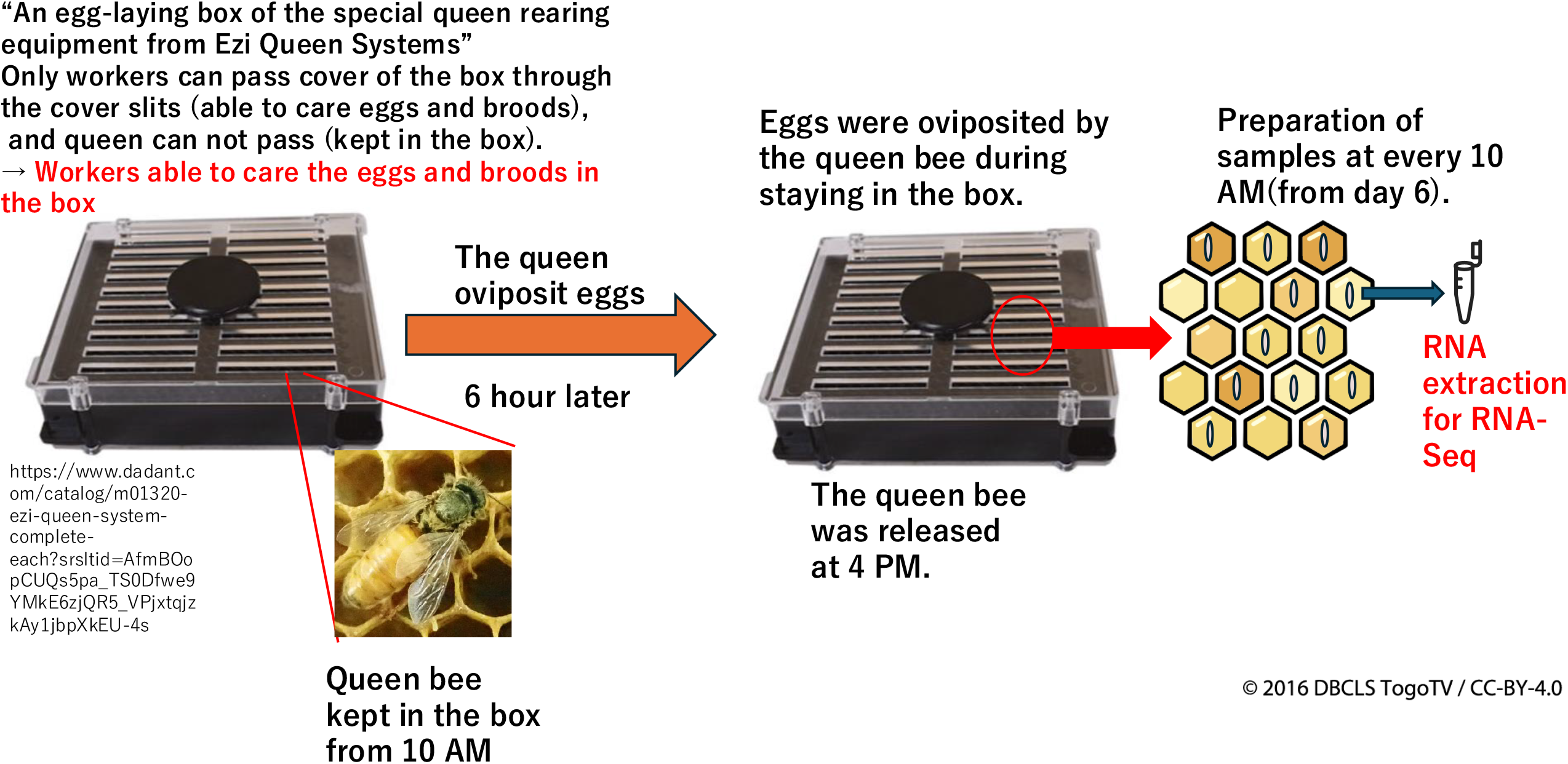
Schematic diagram for the detailed method of sample preparation. Using an egg-laying box, the timing of oviposition by the queen can be controlled (kept in this box from 10 AM to 4 PM). Further, precisely controlled sampling was achieved by sampling at 10 AM . The prepared samples were used for total RNA extraction, and each RNA sample was used for RNA-Seq.

**Fig. 3.**
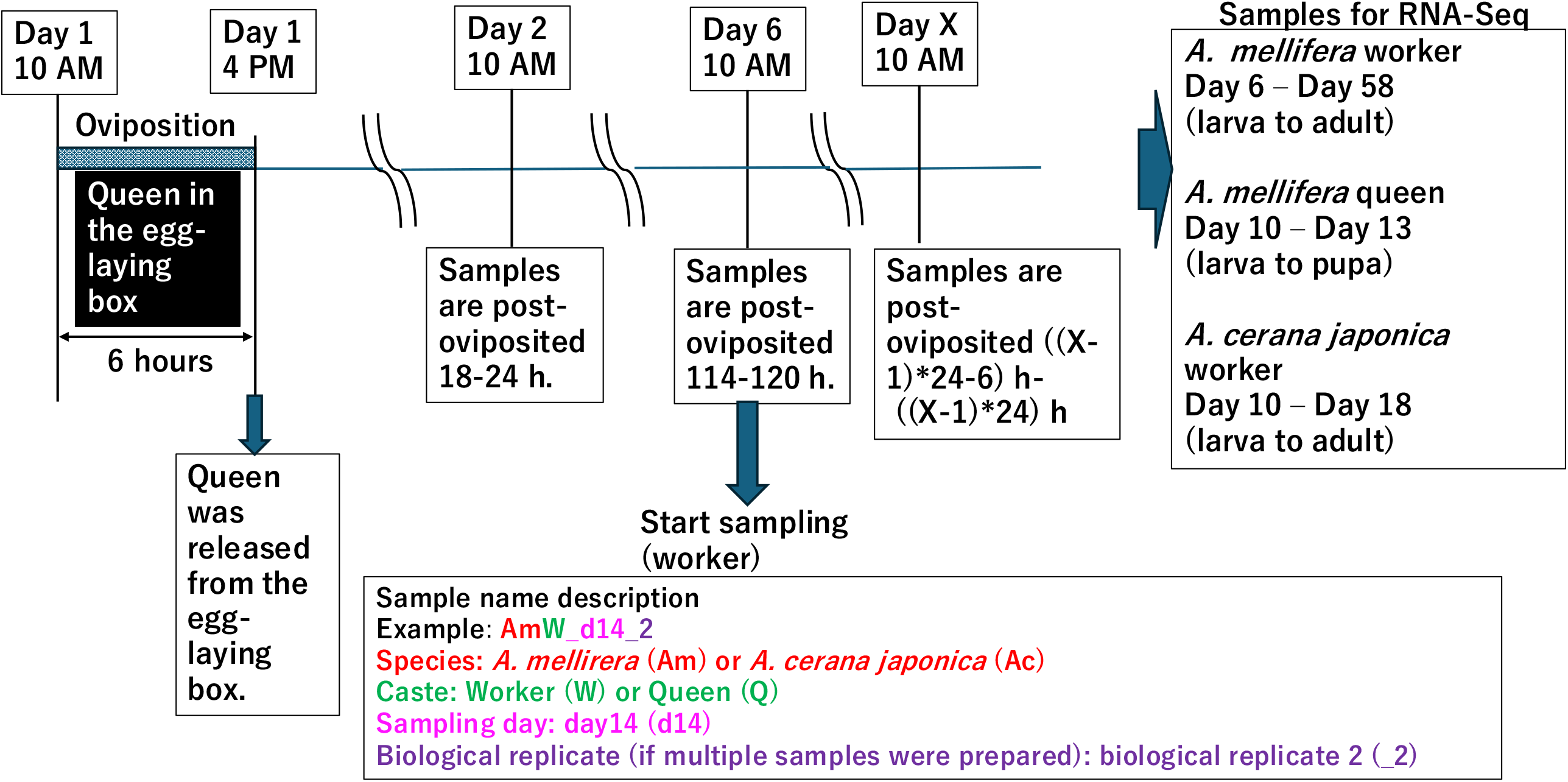
Detailed explanation for the samples in this study. As shown in Fig. 2, one queen was kept in the egg-laying box from 10 AM to 4 PM, for oviposition. For *A. mellifera* workers, sample preparation was started five days afterward (Day 6) at 10 AM. The worker samples were consistently prepared at 10 AM. As shown in this figure, the samples described on Day X were post-oviposited ((X-1)*24-6) hour to (X-1)*24-hour samples. Similarly, *A. mellifera* queen and *A. cerana japonica* worker samples were prepared. In total, *A. mellifera* worker and queen samples were prepared from day 6 to day 58 (larva to adult), and from day 10 to day 13 (adult to larvae), respectively, while *A. cerana japonica* worker samples were prepared from day 10 to day 18 (larvae to adult). Further, the sample name description method was shown at the bottom of part in this figure.

**Fig. 4.**
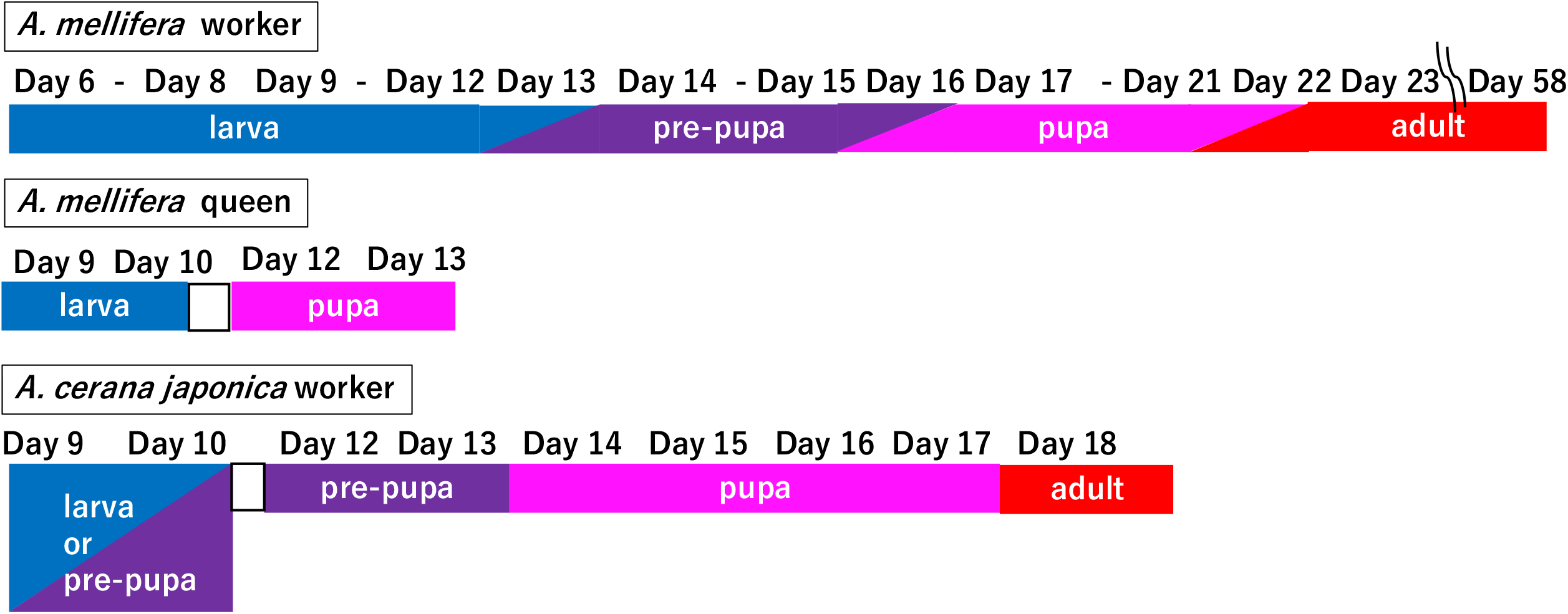
Samples and their developmental stages. Relationships between sample days and developmental stages in this study are shown. For *A. mellifera* samples, the sampling day and developmental stage of each sample were determined as described by Winston (1991)^1^ and based on the morphological features of the samples. *A. mellifera* queen day 11 and *A. cerana japonica* worker day 11 samples were excluded from the sample collections as described in the “Methods” part.

### RNA-extraction

Total RNA was extracted using TRIzol Reagent (Thermo Fisher Scientific, Waltham, MA, USA) according to the product manual. Honey bee samples were homogenized in 2 mL of TRIzol. The extracted RNAs were resuspended in 100 μL of RNase free water and purified using an RNeasy mini kit (Qiagen) according to the product manual. Purified RNA concentrations were quantified using a Nanodrop spectrophotometer (Thermo Fisher Scientific). According to the concentration values and total amounts of extracted RNA, one to five RNA samples (biological replicates) per category were used for RNA-Seq.

### Library preparation and RNA-Seq

cDNA libraries for RNA-Seq were constructed from the extracted RNA samples using the TruSeq Stranded mRNA Library Prep Kit (Illumina, San Diego, CA, USA) following the procedures outlined in the Reference Guide. An Illumina NovaSeq 6000 was used to sequence paired-end reads. Library construction and sequencing were performed by the Macrogen Corp., Japan (Kyoto, Japan).

### RNA-Seq Data analysis

The workflow of the RNA-Seq data analysis used in this study is shown in Fig. 5.

**Fig. 5.**
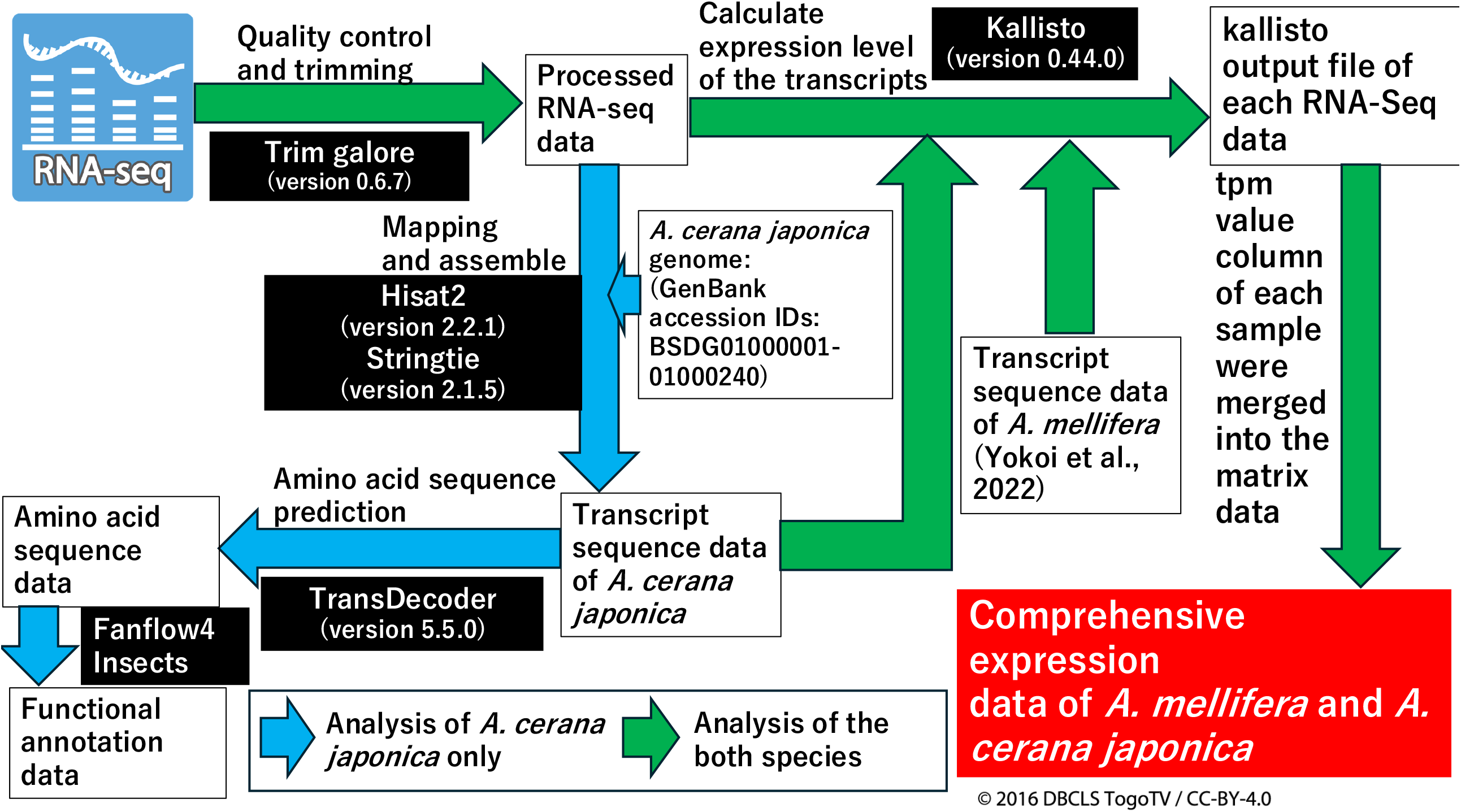
Data analysis workflow in this study. The data analysis workflow, beginning with the raw RNA-Seq data of each sample on the upper left side, is shown. The green arrows indicate the data analysis of both species, *A. mellifera* and *A. cerana japonica*, while light blue arrows indicate the data analysis of *A. cerana japonica* alone. Software names with versions used are shown in the white characters in black boxes.

Removal of adapter sequences, quality control, and trimming of raw RNA-Seq data were performed using Trim Galore (version 0.6.7, https://www.bioinformatics.babraham.ac.uk/projects/trim_galore/) with default settings. Processed RNA-Seq data were used to construct transcriptomic sequences (only *A. cerana japonic*a) and to calculate the expression values of the transcriptome in all prepared samples. To construct transcriptomic sequence data of *A. cerana japonic*a (blue arrows in Fig. 5), the processed RNA-Seq data were mapped to the genome sequences *A. cerana japonic*a using Hisat2 (version 2.2.1)^19^ with the default settings. To construct transcriptome sequence data for each sample, each RNA-Seq data were assembled using the BAM file and gene set data from the genome data using Stringtie^20,21^ (version 2.2.1) with the default settings. The constructed transcriptome sequence data were merged into a single dataset as a GTF file, and the gtf file was converted to the FASTA format using Gffread^22^ (version 0.12.1) as the reference transcriptome sequence data of *A. cerana japonica*. Functional annotations of each transcript were archived using the predicted amino acid sequences (predicted by TransDecoder (version 5.5.0)^23^ with thedefault settings.) by Fanflow4Insects^24^. Protein-level annotations were performed using protein sequence data of comprehensively annotated species (*Homo sapiens, Mus musculus, Caenorhabditis elegans, Drosophila melanogaster, Bombyx mori, Bombus terrestris, Nasonia vitripennis*, and *A. mellifera*) and Unigene data (for details see ref^24^).

Using the generated transcript sequence data of *A. mellifera*^8^ and *A. cerana japonica* (described above) and the RNA-Seq data, the expression levels of these transcripts in all samples of *A. mellifera and A. cerana japonica* were calculated by pseudoalignment of the RNA-seq reads to the reference transcripts with the default settings, using Kallisto^25^ (version 0.44.0). Transcript per million (TPM) values in the Kallisto output files of all samples from the two honey bee species were extracted and merged into single matrix data as comprehensive expression data. All commands and scripts of the data analysis were described in a single text file (see “Code availability” section).

### Hierarchical clustering analysis

To validate the constructed comprehensive expression data, hierarchical clustering analyses (average linkage and Spearman’s rank correlation) were performed using R (version 4.2.3) in Rstudio (version 2023.03.0+386), using the comprehensive expression data of the two species as input. Rscript of the analysis was uploaded in figshare (see “Code availability “ section).

## Results and Discussion

RNA-Seq data and comprehensive Expression data of *A. mellifera* and *A. cerana japonica*

In total, RNA-seq data from eight *A. mellifera* queens, 72 *A. mellifera* workers, and 14 *A. cerana japonica* workers were obtained. In *A. cerana japonica*, reference transcriptome sequence data (47867 transcripts) were constructed using genome sequence data and, 99789 predicted amino acid sequences were constructed using the transcript sequence data. Functional annotation data of the reference transcriptome were constructed using fanflow4insect^23^ based on predicted amino acid sequence data. Next, using the RNA-Seq data and reference transcriptome data of *A. mellifera* and *A. cerana japonica*, expression values of the reference transcriptome in the prepared samples were calculated by pseudoalignment of the RNA-seq reads to the reference transcripts, using Kallisto, and comprehensive expression data of multiple developmental stages were constructed (Fig. 1).

### Data Validation

As described in the “Methods” section, RNA-Seq samples were prepared using an egg-laying box, and a single queen was allowed to oviposit in the egg-laying box (Fig. 2). The queen was introduced and kept in the egg-laying box at 10 AM and released at 4 PM (6 hours) (Fig. 3). Therefore, the eggs in the egg-laying box were laid during the queen’s stay. After several days, samples were prepared at 10 AM, and used for RNA-Seq. Similarly, other samples were consistently prepared at 10 AM. For this sample preparation method, all samples were derived from a single queen and the post-oviposition time of the samples within the same category varied within 6 h. Considering the methods of sample preparation, despite some missing stages in the sample sets of the *A. mellifera queen* and *A. cerana japonica* worker (owing to difficulty in sample preparation), the RNA-Seq data and expression data in this study are reliable as reference expression data.

To validate the reliability of the RNA-Seq and expression data, hierarchical clustering analyses were performed using the expression data of *A. mellifera* and *A. cerana japonica* as input data. The dendrograms of *A. mellifera* and *A. cerana japonica* are shown in Fig. 6 and Fig. 7, respectively. The hierarchical clustering results of *A. mellifera* and *A. cerana japonica* showed that despite several exceptions, two to five biological replicates of the same condition were located in single or closely neighboring clusters, suggesting the reliability of the expression data; further, the clusters containing samples from relatively close sampling days were formed.

**Fig. 6.**
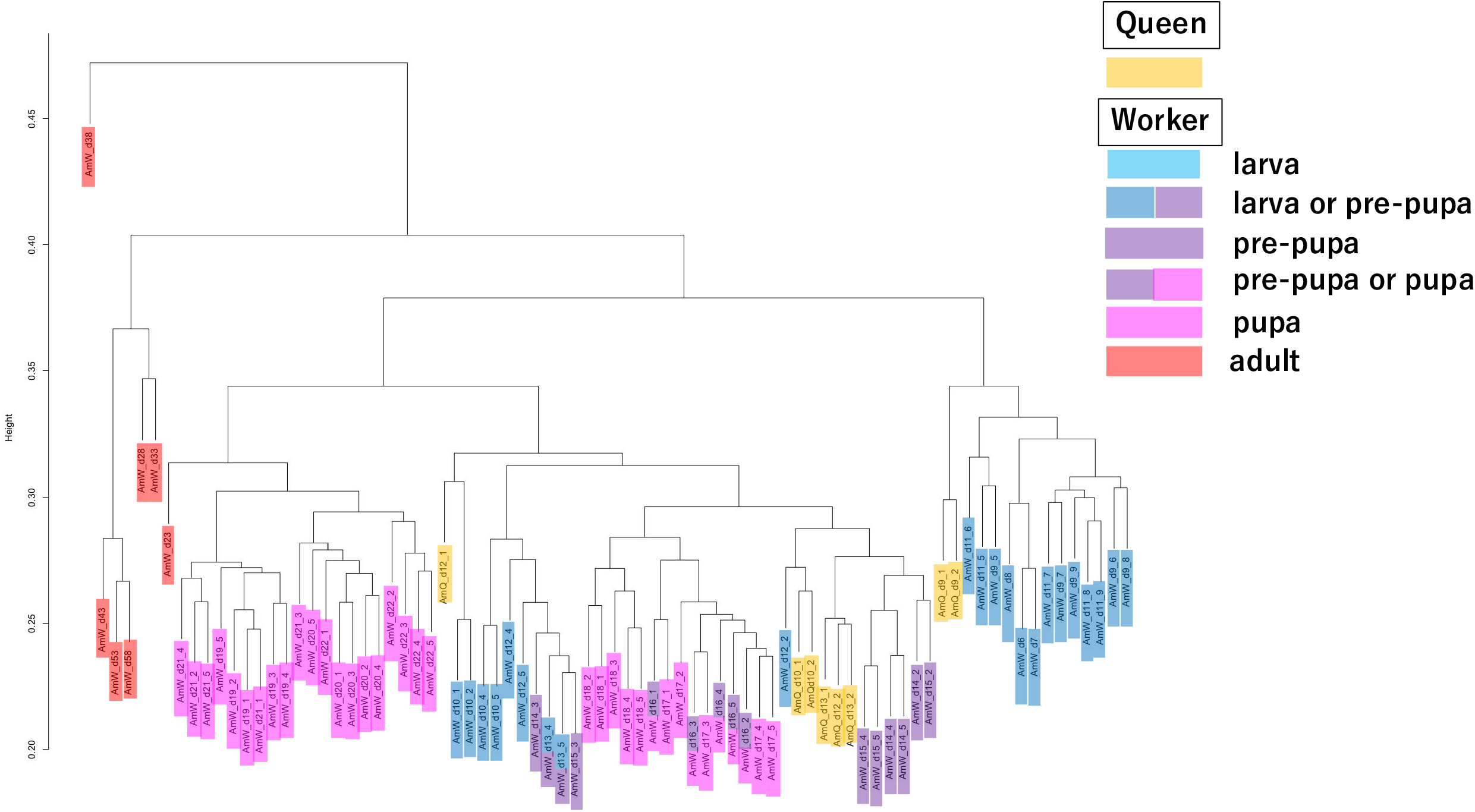
Dendrogram of hierarchical clustering analysis using *A. mellifera* expression data. Hierarchical clustering analysis was performed using *A. mellifera* expression data as input. The dendrogram was drawn based on the result of the hierarchical clustering analysis. In the dendrogram, queen samples were colored yellow. In worker samples, larva, pre-pupa, pupa, and adult samples are colored light blue, blue, purple, pink, and red, respectively. The sample category containing two stages are indicated in two colors (e.g., samples containing both larva or pre-pupa stage are colored in blue and purple). The manner of coloring is the same as that in Fig. 4. The biological replicate numbers of some sample categories do not begin with “1” and are not sequential (See “Methods” part.).

**Fig. 7.**
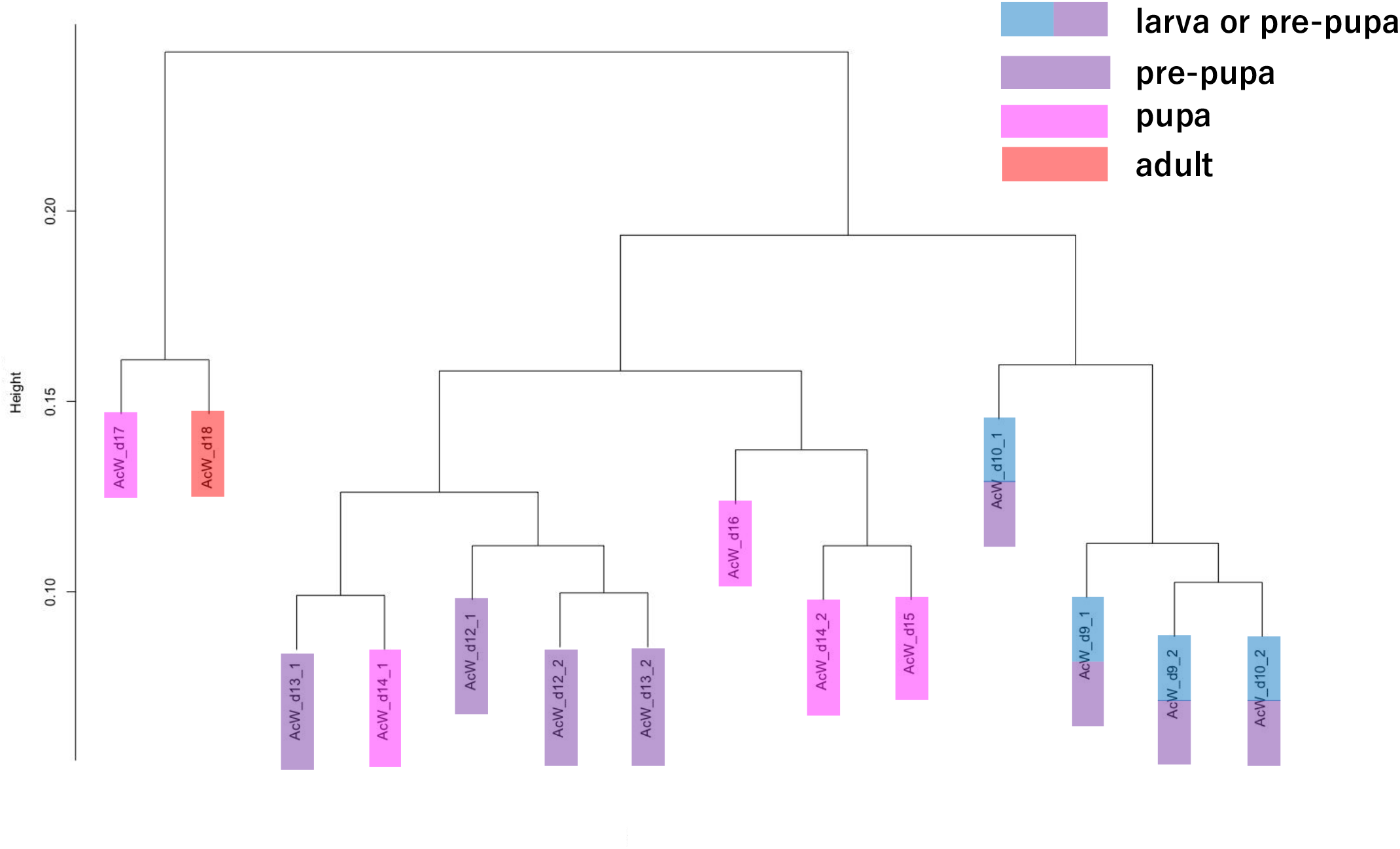
Dendrogram of hierarchical clustering analysis using *A. cerana japonica* expression data. Hierarchical clustering analysis was performed by using *A. cerana japonica* expression data as input. The manner of the sample coloring is the same as that in Fig. 4.

In *A. mellifera* clustering results (Fig. 6), all *A. mellifera* worker adults (colored in red), except AmW_d23 (immediately after becoming adults), were first separated, revealing that *A. mellifera* adult workers have different transcriptome profiles compared to the their pupal or larval stage samples. Adult workers perform multiple tasks in colonies such as caring for broods and foraging. Considering these traits, the separation of the transcriptome profiles of *A. mellifera* adult workers from other samples is justified. Some of the larval worker samples (AmW_d6-9 and d11), and AmQ_d9 samples (larvae) formed one cluster (on the right side). This cluster may reflect the transcriptome profile of larval stages. The other clusters included the AmW_d10 and AmW_d12 samples (larvae); AmW_d13 to d18 (pre-pupa to pupa); and AmQ_d10, d12, and d13 (larvae and pupa). These clusters may reflect the transcriptome profiles of the prepupal to early pupal stages. We assumed that the AmW_d10 samples developed faster than usual^1^, although they were from the late larval stage, while AmW_d12 and AmQ_d10 samples were from the late larval stage, but their transcriptome profiles were shifted to those in the pre-pupal and pupal stages prior to the morphological changes from larva to pre-pupa. The AmQ_d12_1 sample was located at an isolated position, possibly because the developmental time differed from that of the other queen samples. Generally, the developmental duration of the queen (from egg to adult) is shorter than that of the worker^1^. AmW_d19-d23 samples (pupae and adults) formed another cluster, possibly reflecting the transcriptome profile of emergence, which drastically changes the morphological traits. The inclusion of the AmW_d19 and d20 samples (mid-pupal stage) in the cluster may be attributed to the shift in transcriptome profiles from AmW_d18 samples to these samples occurring prior to the morphological changes during emergence. In *A. cerana japonica*, all samples were divided into three clusters (Fig. 7). One cluster included the AcW_d17 and AcW_d18 samples (pupae and adults), which may reflect the transcriptome profiles of the early adult stage or emergence. The other cluster included AcW_d12–d16 samples (pre-pupa and pupa). The third cluster included the AcW_d9 and d10 samples (larvae and pre-pupa). These clusters may reflect transcriptome profiles from the pre-pupa to pupa and the larva or early pre-pupa stages, respectively. These results reflect the biological traits of the *A. cerana japonica* developmental stages. In conclusion, both clustering results suggest that the transcriptome data in this study are reliable and can be used as comprehensive reference expression data (reference transcriptome data).

## Conclusion

RNA-seq data from eight *A. mellifera* queens, 72 *A. mellifera* workers, and 14 *A. cerana japonica* workers were obtained. The validation results of the RNA-Seq and comprehensive expression data described below indicate the reliability of the reference transcriptome and expression data, which can contribute to honey bee research (e.g., determining target genes to create strains with beneficial traits for the beekeeping industry by genome editing) or evolutionary and comparative biology between insect species or social insect species at the genetic and molecular levels. These comprehensive expression data are the first public reference expression data (reference transcriptome data) of *A. mellifera* and *A. cerana japonica* samples derived from a single queen.

## Code Availability

The code files for transcriptome analysis shown in Fig. 5 and hierarchical clustering analysis using R as described in “Technical Validation” section are available in figshare^33^.

## Data Records

The raw RNA-seq data in this study were deposited in the Sequence Read Archive (SRA, ID: DRA016719 and DRA014424)^26,27^. The BioProject, BioSample, and SRA accession IDs for each RNA-seq sample are described in the table file uploaded to figshare^28^. The assembled transcript data in gtf and FASTA files, the predicted amino acid sequence FASTA file, and the functional annotation file of the *A. cerana japonica* transcripts are available in figsahre^29^. The TPM value matrix data of all transcripts in all samples of *A. mellifera*^30^ and *A. cerana japonica* were prepared^31^ and the Kallisto output files for the construction of matrix data were uploaded to figshare^32^.

## Acknowledgements

This work was supported by the Center of Innovation for Bio-Digital Transformation (BioDX) and an open innovation platform for industry-academia co-creation (COI-NEXT) of JST (COI-NEXT, JPMJPF2010) to K.Y., M.H., S.K., A.J., H.B., and K.K. Parts of Figs. 1, 2, and 5 were drawn using illustrations from TogoTV (© 2016 DBCLS TogoTV, CC-BY-4.0 https://creativecommons.org/licenses/by/4.0/deed.ja).

We would like to thank Editage (www.editage.jp) for English language editing.

We express thankfulness to Dr. Shotaro Mine at Institute of Agrobiological Sciences NARO for the preparations of A. mellifera queen photographs used in Figure 2.

## Author contributions

K.Y., M.H., H.B. and K.K. conceived the study. M. H., S. K., T. M., M. Y., M. O. H., S. M., and K. K. prepared the RNA samples from *A. mellifera* and *A. cerana japonica* for RNA-seq. K.Y., A.J., and H.B. performed the bioinformatics data analysis and data registration. K.Y wrote the original draft of the manuscript. All authors reviewed and edited the draft of the manuscript. All the authors have read and agreed to the published version of this manuscript.

